# Fetal reversion from diverse lineages sustains the intestinal stem cell pool and confers stress resilience

**DOI:** 10.1101/2025.07.08.663108

**Authors:** Sakura Kirino, Fumiya Uefune, Kensuke Miyake, Nobuhiko Ogasawara, Sakurako Kobayashi, Satoshi Watanabe, Yui Hiraguri, Go Ito, Keiichi Akahoshi, Daisuke Ban, Johan H. van Es, Hans Clevers, Mamoru Watanabe, Ryuichi Okamoto, Shiro Yui

## Abstract

Plasticity is a central mechanism underlying the robust regenerative capacity of the intestinal epithelium. Two major plasticity have been described: spatial plasticity, in which differentiated cells revert to crypt base columnar cells (CBCs), and fetal reversion into revival stem cells (revSCs). However, the relationship among these two stem cell populations and differentiated cells remains to be clarified.

Here, we demonstrated the bidirectional interconversion between CBCs and revSCs. Using lineage tracing, injury models and villus culture, we show that absorptive enterocytes can reprogram into revSCs and regenerate CBCs. These findings position fetal reversion as an entry point to spatial plasticity, establishing a regenerative hierarchy where CBCs, revSCs, and enterocytes collectively orchestrate intestinal repair. Furthermore, we identified revSCs as a highly stress-tolerant stem cell population, whose emergence is crucial for preserving the stem cell pool.

Our results establish fetal reversion as a cellular escape mechanism safeguarding epithelial regeneration under inflammatory conditions.

## INTRODUCTION

The intestinal epithelium exhibits a remarkable capacity for regeneration following injury. Crypt base columnar stem cells (CBCs), located at the base of the intestinal crypts of Lieberkühn governs the homeostasis in normal intestinal epithelium^1^. These intestinal stem cells are defined by the expression of markers such as *Lgr5*^2^ and *Olfm4*^2, 3^. As they migrate from the crypt base toward the villus tip, CBCs differentiate into the various cell types that comprise the intestinal epithelium, including absorptive enterocytes (approximately 90% of epithelial cells) and secretory lineages such as goblet, Paneth, enteroendocrine, and tuft cells^4^.

In response to epithelial injury, various differentiated cell types have been shown to contribute to the restoration of the CBC pool^5, 6, 7, 8, 9^. This process, termed *spatial plasticity*^5, 10, 11^, is thought to involve positional shifts of mature cells into the stem cell niche, where they reacquire stemness after loss of CBCs. More recently, *revival stem cells* (revSCs)—a population transcriptionally reminiscent of fetal progenitors—have been identified as key players in tissue regeneration^12^. This phenomenon, now referred to as *fetal reversion*^12, 13, 14^, is under intense investigation in the context of inflammatory bowel diseases (IBD)^15,16^ and colorectal cancer^17, 18^. The intestinal epithelium is thus emerging as a therapeutic target beyond immune modulation^19, 20^ and understanding the molecular and cellular mechanisms of regeneration is a crucial in translational medicine. Despite recent advances, the relationship between spatial plasticity and fetal reversion remains to be clarified. In particular, how CBCs, revSCs, and differentiated lineages interrelate within regenerative hierarchies is unclear. Moreover, the biological advantages conferred by such dynamic cellular transitions during regeneration remain poorly understood.

In this study, we demonstrate bidirectional conversion between CBCs and revSCs, confirming the ability of CBCs to generate revSCs, and vise verse from revSCs to CBCs. In addition, flow cytometry, pseudotime trajectory analyses, and lineage tracing experiments revealed that absorptive enterocytes contribute to the revSC pool. Importantly, enterocyte-derived revSCs could generate CBCs in lineage tracing experiments of villus culture coupled with organoid transplantation model^21, 22^, organoid replating assay^14^, and 5-FU enteritis model^23, 24^, providing evidence that fetal reversion serves as an entry point to spatial plasticity. Notably, while classical organoid systems have long been established exclusively from crypt-resident stem cells^25, 26^, our findings challenge this longstanding paradigm by demonstrating that mutation-free villus-derived cells can indeed establish organoids via fetal reversion, revealing remarkable regenerative flexibility of the adult intestinal epithelium. Finally, in vitro 5-FU treatment revealed a remarkable potential of revSCs to resist for cellular stress.

Together, our findings propose a conceptual model in which revSCs primarily arise from CBCs and can also be supplied by absorptive enterocytes, integrating spatial plasticity and fetal reversion into a unified regenerative program. Functionally, revSCs constitute bona fide stem cells equipped with enhanced stress resistance, allowing for epithelial repair and stem cell preservation under inflammatory conditions.

## RESULTS

### Single cell RNA sequence estimates CBCs and absorptive enterocyte as an origin of revival stem cells

To identify candidate cell populations with the potential to undergo fetal reversion and acquire revSC identity, we employed a 3D organoid culture system derived from mouse proximal small intestine. We compared two conditions: standard budding organoids cultured in Matrigel (Matrigel organoids)^25^, which recapitulate the homeostatic epithelium, and revSC rich spherical organoids cultured in Collagen type I (Collagen organoids), previously shown to induce YAP/TAZ-dependent fetal reprogramming and inflammatory epithelial phenotypes ^14^.

Single-cell RNA sequencing (scRNA-seq) of these two organoids revealed eight transcriptionally distinct epithelial clusters (Figure 1A, B, Figure S1A). Matrigel organoids contained diverse intestinal cell types, including CBCs cells and differentiated lineages, whereas Collagen organoids displayed a transcriptionally homogeneous profile dominated by revSC-like cells (Figure 1B, Figure S1A). *Ly6a*, a canonical revSC marker^13, 14^, was exclusively expressed in cluster 1, corresponding to Collagen organoid–derived cells (Figure 1C, Figure S1B). In addition, a previously reported revSC signature (*Ly6a, Clu, Mif, Anxa1,* and *Anxa3*)^12, 14^ was highly enriched in this cluster (Figure 1C). In contrast, the CBC signature (*Lgr5, Olfm4, Smoc2,* and *Ascl2*)^2, 27^ was distributed across clusters 2, 3, and 4, reflecting the stem cell rich nature of Matrigel organoid cultures^28, 29^ (Figure 1B,C).

**Figure 1.**
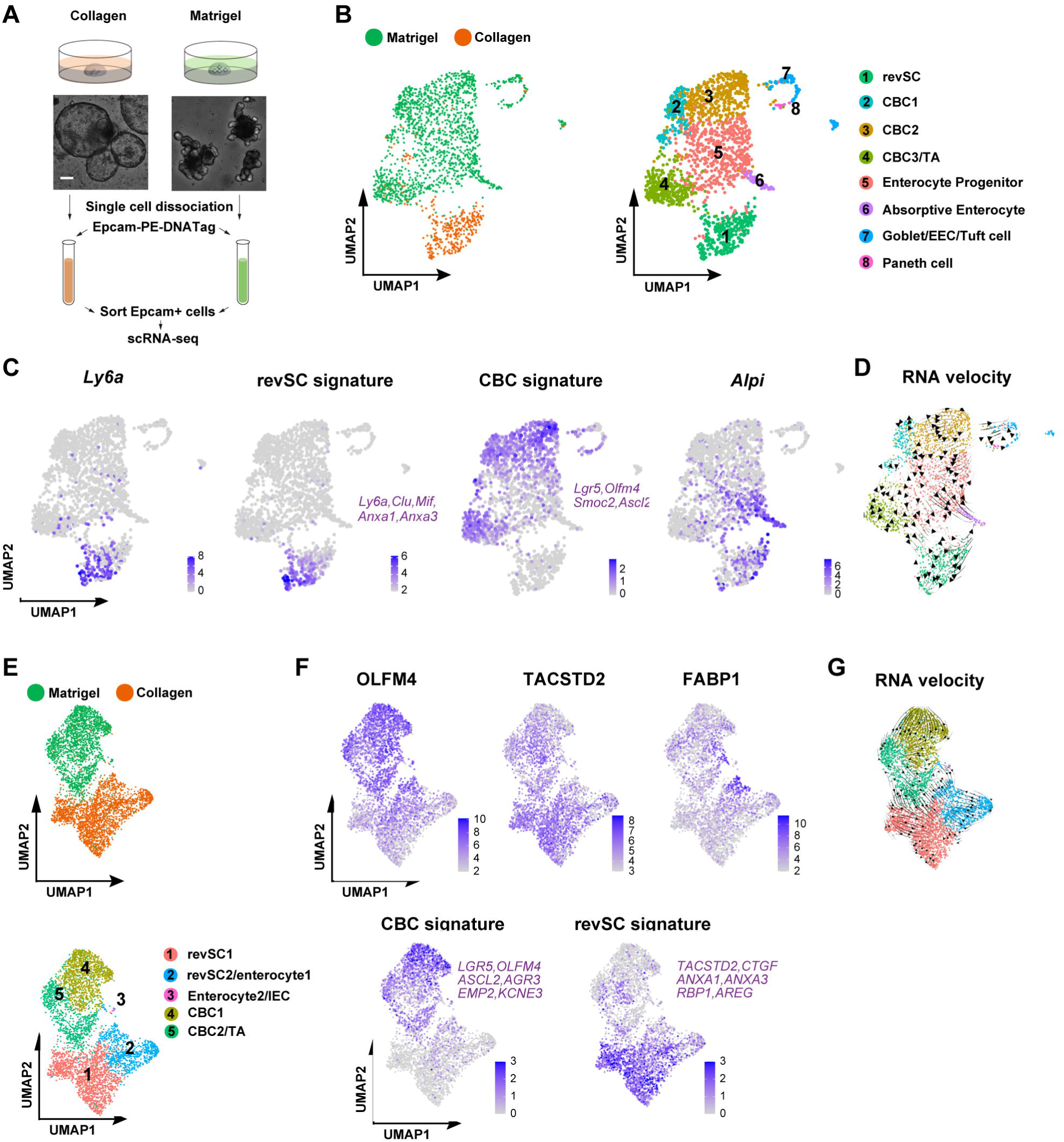
Single cell RNA sequence estimates CBCs and absorptive enterocyte as an origin of revival stem cells. (A) Schematic overview of the single-cell RNA sequencing workflow. Mouse organoids in Matrigel and Collagen were dissociated, and DNA-barcoded epithelial cells (Epcam+) were isolated by fluorescence-activated cell sorting (FACS). Scale bar: 200µm. (B) UMAP visualization of epithelial cells from mouse organoids, colored by sample (left) and by annotated cell type (right). Clusters were assigned based on known and top marker gene expression. (C) Feature plots of *Ly6a*, revSC signature, CBC signature, and *Alpi*, showing spatial expression patterns or module scores projected onto the UMAP embedding. (D) RNA velocity analysis of mouse organoids by scVelo package. Arrows indicate the direction and speed of cell state transitions. (E) UMAP visualization of epithelial cells from Human organoids, colored by sample (top) and by annotated cell type (bottom). Clusters were assigned based on known and top marker gene expression. (F) Feature plots of *OLFM4*, *TACSTD2*, *FABP1*, CBC signature, and revSC signature, showing spatial expression patterns or module scores projected onto the UMAP embedding. (G) RNA velocity analysis of human organoids by scVelo package. Arrows indicate the direction and speed of cell state transitions.

*Alkaline phosphatase, intestinal (Alpi)*, a marker of absorptive enterocytes^5^, was predominantly detected in clusters 5 and 6 (Figure 1C). To explore lineage dynamics, we performed RNA velocity analysis using scVelo^30^. The inferred trajectory flow suggested that revSCs emerged from both *Lgr5+* CBC/transit-amplifying (TA) populations and Alpi+ enterocyte lineage (Figure 1D), indicating that reversion process has two distinct routes.

We next validated these findings in human small intestinal organoids using scRNA-seq. In contrast to mouse Collagen organoids, which predominantly consisted of a single undifferentiated cell population, human Collagen organoids exhibited two distinct clusters, annotated as cluster 1 and 2 (Figure 1E, Figure S1C). In human Matrigel organoids, we identified a limited number of transcriptionally distinct clusters, classified as CBCs (cluster 4), CBC/TA cells (cluster 5), and a small minor cluster (cluster 3), which likely represents a subset of absorptive enterocytes among intestinal epithelial cells (IECs) (Figure 1E, Figure S1C).

Feature plots visualization revealed a reciprocal expression pattern of *OLFM4* and *TACSTD2*, established markers of the fetal intestinal state^15^ indicating that Matrigel organoids predominantly exhibit a mature, adult epithelial phenotype, whereas Collagen organoids acquire a revSC-like phenotype consistent with fetal reversion (Figure 1F). Notably, *FABP1* expression was enriched in a subset of clusters 2 and 5, consistent with their assignment to the absorptive enterocyte lineage (Figure 1F). Dot plot analysis further delineated gene expression signatures specific to each cluster (Figure S1C), enabling the identification of canonical markers for CBCs (*LGR5, ASCL2, OLFM4, AGR3, EMP2,* and *KCNE3*) and for revSCs (*CCN2, TACSTD2, ANXA1, ANXA3, RBP1,* and *AREG*)^12, 14, 31^. These marker gene sets facilitated not only the distinction between Matrigel and Collagen organoids but also revealed phenotypic heterogeneity within the Collagen clusters: cluster 1 representing a putative revSC-dominant population, and cluster 2 representing a mixed revSC/enterocyte state (Figure 1F, Figure S1C) ^32, 33^. RNA velocity analysis using scVelo further revealed two major lineage trajectories: one progressing from CBC/TA to revSCs, and the other from absorptive enterocytes lineage to revSCs, mirroring the dual bifurcation observed in the mouse model (Figure 1G). Importantly, all six revSC signature genes identified in human organoids were previously reported to be upregulated in actively inflamed human intestinal mucosa^34^, underscoring the relevance of revSCs as an inflammation-associated epithelial state.

Together, these results define revSCs as a transient, niche-dependent epithelial state arising from both CBCs and absorptive enterocytes under specific ECM conditions, reflecting a plastic response of both the mouse and the human intestinal epithelium to microenvironmental cues.

### CBCs serve as a major source of revSCs, and the two stem cell populations are mutually interconvertible

To experimentally define the cellular sources of revSCs, we employed a fetal reversion model using Collagen organoid cultures. Based on the scRNA-seq findings suggesting that *Lgr5*+ populations harbor the potential to become revSCs, we performed a reconstitution assay using *Lgr5^EGFP-IRES-CreERT2/+^* mice ^2^ to evaluate fetal reversion capacity among distinct cell populations under homeostatic conditions, stratified by their stemness (Figure 2A). Flow cytometric analysis enabled separation of epithelial cells into GFPhigh CBCs, GFPlow progenitors, and GFPneg differentiated cells ^25^ (Figure 2B). Sorted populations were embedded in either Matrigel or Collagen and cultured in the same reconstitution medium. Organoid formation efficiency (OFE) was significantly higher in the GFPhigh CBC population in both matrices (Figure 2C, 2D), indicating high reversion potential. These findings are consistent with the predictions from prior computational modeling^31, 35^. Notably, Matrigel organoids maintained *Lgr5* expression and did not express *Ly6a*, whereas Collagen organoids showed broad *Ly6a* expression and lacked *Lgr5*, consistent with a revSC phenotype (Figure 2E). Interestingly, both GFPlow and GFPneg populations exhibited significantly higher OFE in Collagen compared to Matrigel (Figure 2D), suggesting that progenitor and differentiated cells retain fetal reversion potential, which can be induced in a niche-dependent manner.

**Figure 2.**
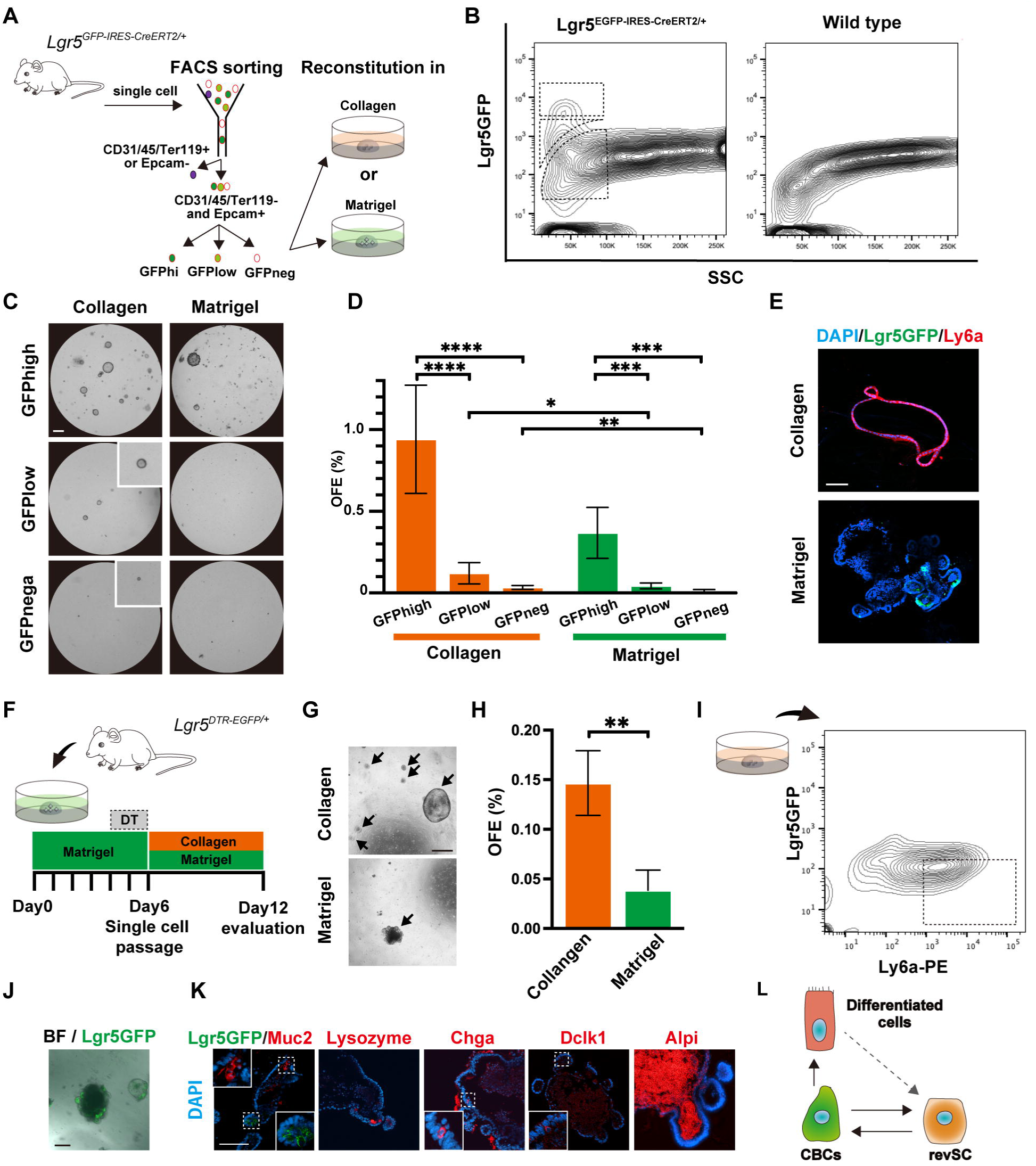
CBCs serve as a major source of revSCs, and the two stem cell populations are mutually interconvertible. (A) Reconstitution assay using small intestinal epithelial cells (CD31-, CD45-, Ter119-, EpCAM+) isolated from *Lgr5^EGFP-IRES-CreERT2/+^* mice. Cells were classified into GFPhigh, GFPlow, and GFPneg populations by FACS. Sorted cells were suspended in Matrigel or Collagen matrices and assessed for organoid formation efficiency (OFE). (B) Lgr5GFP+cells from *Lgr5^EGFP-IRES-CreERT2/+^* mouse (left) and wild-type mice (right). Representative gating strategies are shown for each population. (C) Representative images of organoid cultures 6 days after seeding equal numbers of cells. Scale bar: 500µm. (D) Comparison of OFE under each culture condition. Sorted cells were cultured in either Collagen or Matrigel, and OFE was quantified. Data are presented as mean ± SD. Statistical significance among different populations within the same matrix condition (Collagen or Matrigel) was assessed using one-way ANOVA (n = 5; Collagen: F(2,12) = 32.90, ****P < 0.0001; Matrigel: F(2,12) = 23.67, \*\*\**P* < 0.001), followed by Tukey’s multiple comparisons test. For pairwise comparisons between Collagen and Matrigel within the same GFPlow or GFPneg population, two-tailed Student’s *t*-test were performed (n = 5; \*\**P* < 0.01 and \**P* < 0.05, respectively). (E) Immunohistological analysis of reconstituted *Lgr5^EGFP-IRES-CreERT2/+^* organoids (shown in Figure 2C). Both samples were stained with anti-Ly6a antibody (red). Collagen organoids exhibited a revSC phenotype (high Ly6a expression and absence of LGR5), while Matrigel organoids contained cells with a CBC phenotype (high LGR5 expression but negative for Ly6a). Images are counterstained with DAPI (blue). Scale bar: 100 µm. (F) Comparison of OFE after CBC deletion and passaging. CBCs were ablated in Matrigel organoid cultures by treatment with diphtheria toxin (DT; 0.05µM) from day 4 to day 6, followed by passaging into Collagen or Matrigel. Organoid formation efficiency was subsequently evaluated on day 12. (G) Representative images of organoid cultures 6 days after seeding equal numbers of cells. Arrow indicates organoids. Scale bar: 500 µm. (H) Comparison of OFE in Collagen or Matrigel culture after CBC deletion and dissociation into single cells. Organoids were formed more efficiently under Collagen culture conditions. Data are presented as mean ± SD (n = 6, two-tailed Student’s *t*-test, \**P* < 0.05). (I) Reconstitution assay from revSCs to CBCs. Organoids derived from *Lgr5^DTR-EGFP/+^* mice were cultured in Collagen. Ly6a+/Lgr5− revSCs were isolated by FACS and seeded into Matrigel. (J) Reconstituted budding organoids derived from the revSC population. Representative immunofluorescence (IF) images show reappearance of Lgr5− GFP+ CBCs at the crypt domain. Scale bar: 100 µm. (K) Immunofluorescence images of cell-specific markers of the reconstituted organoids shown in (J). Boxed regions are enlarged at the periphery of the panel. Reappearance of Lgr5+ CBCs (green), Muc2+ goblet cells (red), Lyz+ Paneth cells (red), Chga+ enteroendocrine cells (EECs; red), Dclk1+ Tuft cells (red), and Alpi+ absorptive enterocytes (red). Images are counterstained with DAPI (blue). Scale bar: 100 µm. (L) A schematic presentation to explain 1) CBCs exhibit the highest fetal reversion potential, 2) differentiated epithelial cells can be reprogrammed into revSCs, and 3) the two stem cell populations are mutually interconvertible.

Given the mosaic expression of GFP in *Lgr5^EGFP-IRES-CreERT2^* mice due to gene silencing^36^ identification of Lgr5-negative populations remains ambiguous. To solve this issue, we utilized *Lgr5^DTR-EGFP^ mice*^37^ for CBC deletion. Organoids were established and subjected to diphtheria toxin (DT) treatment between day 4 and day 6 for CBCs deletion (Figure 2F). Single-cell suspensions were replated in either Matrigel or Collagen matrices, and OFE in each condition was quantified. Organoid formation was significantly enhanced in Collagen culture (Figure 2G, H), confirming that fetal reversion is promoted even in the absence of CBCs in niche-dependent manner. Of note, minimal organoids formation in Matrigel post-CBC deletion is consistent with previous reports of organoid regeneration from reserve populations^5, 37^. Next, to examine whether revSCs can give rise to CBCs oppositely, we established Collagen organoids from *Lgr5^DTR-EGFP^* mice, sorted Ly6a+/Lgr5− revSCs population, and replated them into Matrigel (Figure 2I). As expected, budding organoids containing Lgr5GFP+ CBCs emerged (Figure 2J), and immunofluorescence analysis confirmed the presence of differentiated intestinal cell types including Muc2+ goblet cells, Lyz+ Paneth cells, Chga+ enteroendocrine cells, Dclk1+ tuft cells, and Alpi+ absorptive enterocytes (Figure 2K). These findings validate the capacity of revSCs to generate functional CBCs and support the notion that revSCs represent bona fide stem cells.

Together, our findings demonstrate that both progenitors and differentiated cells retain the capacity for fetal reversion, underscoring the cooperative interplay between fetal reversion and spatial plasticity in orchestrating epithelial regeneration. Also, CBCs display a particularly high capacity for fetal reversion, highlighting their central role in initiating regenerative transitions. Furthermore, revSCs and CBCs exhibit mutual interconvertibility, representing two distinct but plastic stem cell states within the intestinal epithelium.

### Lineage tracing experiments revealed fetal reversion potential of differentiated enterocytes in vitro

To understand the entire process of fetal reversion, contribution of non-CBCs is not negligible as shown above. Accumulating evidence has also demonstrated that various differentiated epithelial cell types can contribute to stem cell pool restoration via spatial plasticity^5, 6, 7, 9, 38^. RNA velocity analysis of our scRNA-seq data predicts a directional shift from absorptive enterocytes toward revSC-like states. To experimentally confirm this lineage flexibility, we performed in vitro lineage tracing using *Alpi^CreER/+^; Rosa^tdTomato/+^* mice. Matrigel organoids established from proximal small intestine were labeled with tdTomato following 4-hydroxytamoxifen (4-OHT) treatment, thereby marking Alpi+ enterocytes localized to the villus domain and their progeny (Figure 3A, 3B). These organoids were serially passaged in Matrigel or Collagen. While tdTomato+ cells gradually disappeared during passaging in Matrigel, they were stably retained in Collagen organoids, suggesting survival or expansion of lineage-traced cells (Figure 3B, 3C). Flow cytometric analysis further revealed that these tdTomato+ cells in Collagen organoids expressed Ly6a, a hallmark of revSCs, at comparable levels to tdTomato-cells (Figure 3D), indicating that revSCs derived from Alpi+ enterocytes are molecularly indistinguishable from other sources of revSCs. To further examine whether these revSCs could reconstitute homeostatic intestinal epithelium, tdTomato+ Collagen organoids were replated into Matrigel. As a result, budding organoids re-emerged, and the entire organoid structure, including the crypt domain were labeled tdTomato+ (Figure 3B). Immunofluorescence analysis confirmed the presence of multiple lineages, including Olfm4+ CBCs, Muc2+ goblet cells, Lyz+ Paneth cells, Chga+ enteroendocrine cells, Dclk1+ tuft cells, and Alpi+ absorptive enterocytes (Figure 3E), indicating that enterocyte-derived revSCs restored a fully functional intestinal epithelial architecture in vitro.

**Figure 3.**
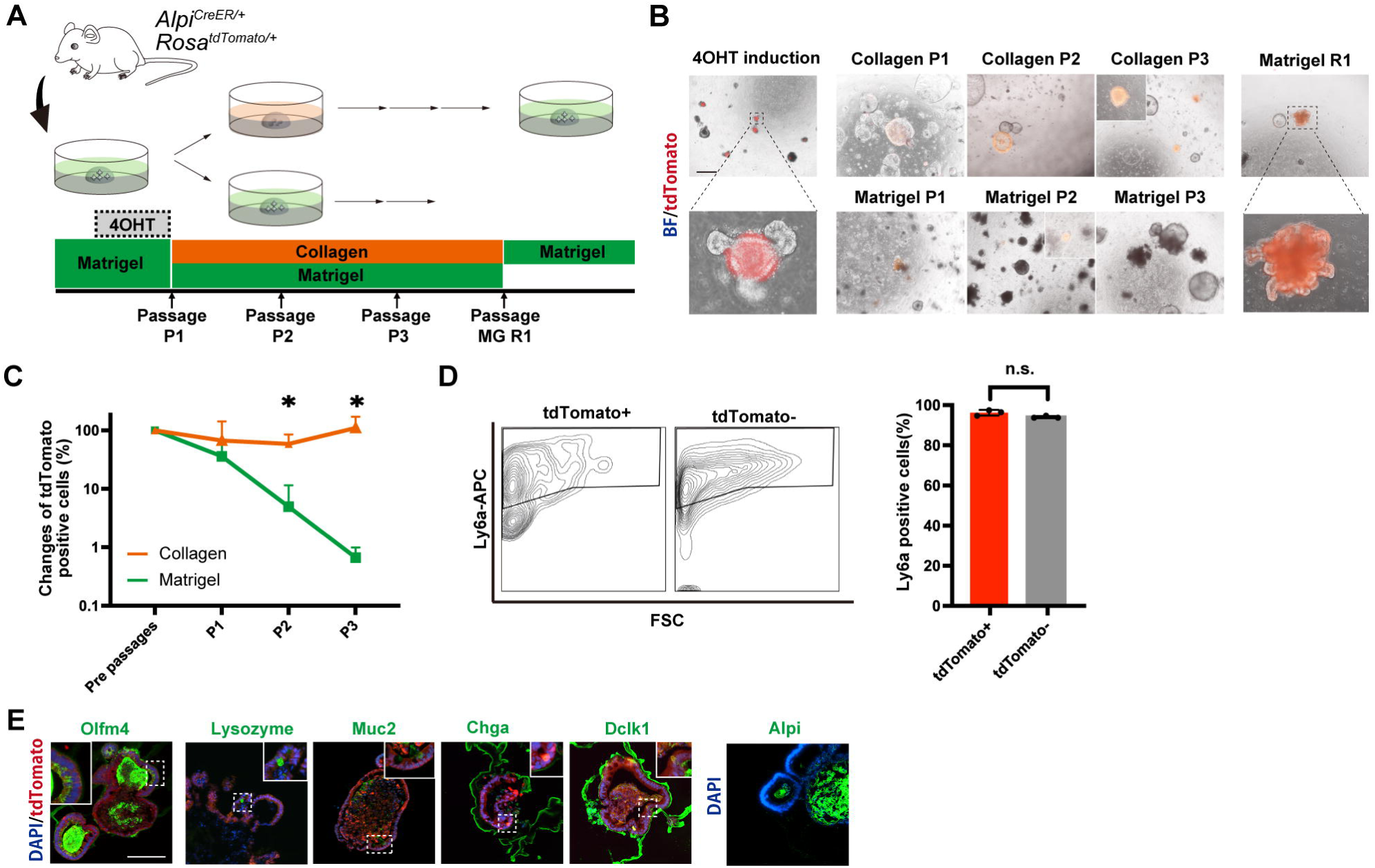
Lineage tracing experiments revealed fetal reversion potential of differentiated enterocytes in vitro. (A) Lineage tracing of Alpi+ absorptive enterocytes in vitro. Organoids were established from *Alpi^CreER/+^; Rosa^tdTomato/+^* mice and treated with 4-OHT to label Alpi+ absorptive enterocytes with tdTomato. After labeling, organoids were serially passaged three times in Collagen or Matrigel. Collagen organoids were subsequently replated into Matrigel. (B) Representative fluorescence images of tdTomato-labeled cells (red) were acquired on day 6 of each passage. Boxed regions are enlarged below. Shortly after labeling, tdTomato+ cells were restricted to the villus domain, which were lost during passages. In collagen organoids however, they remained detectable during passages. Upon replating into Matrigel, tdTomato+ cells reconstituted entire organoids, including the crypt domain. Scale bar: 500 µm. (C) Quantitative analysis of tdTomato+ cells by FACS. Data are presented as mean ± SD. Collagen organoids retained significantly more numbers of tdTomato+ cells than those in Matrigel at P2 and P3 (n = 3, two-tailed Student’s *t*-test, \**P* < 0.05). (D) Comparison of Ly6a+ cell frequency between tdTomato+ and tdTomato-populations. Representative FACS plots (left) and quantification (right) are shown. No significant difference was observed (n=3, two-tailed Student’s *t*-test, *P* = n.s.) (E) Immunofluorescence images of cell-specific markers of the Matrigel-R1 organoid shown in (B), which is entirely composed of tdTomato+ cells. Boxed regions are enlarged at the periphery of the panel. The organoid is composed of Olfm4+ CBCs (green), Lyz+ Paneth cells (green), Muc2+ goblet cells (green), Chga+ EECs (green), Dclk1+ Tuft cells, and Alpi+ absorptive enterocytes (green). Images are counterstained with DAPI (blue). Scale bar: 100 µm.

These findings provide direct evidence that differentiated absorptive enterocytes can undergo fetal reversion in niche-dependent manner and subsequently regenerate functional CBCs, implying a lineage plasticity that contributes to epithelial regeneration.

### Fetal reversion enables villus-derived cells to regenerate tissue without oncogenic mutations

To determine whether the fetal reversion and subsequent conversion to CBCs observed in Alpi+ enterocytes in vitro could also occur under more physiological conditions, we performed primary Collagen culture of villus tissues containing tdTomato labeled Alpi+ cells, followed by in vitro organoid expansion and subsequent transplantation experiments using a DSS-induced colitis model^21, 22^. *Alpi^CreER/+^; Rosa^tdTomato/+^* mice were treated with tamoxifen to label Alpi+ enterocytes and their progeny (Figure S2A). Two days later, villi were isolated from the proximal small intestine as previously described^5, 39, 40^, and embedded in Collagen or Matrigel matrices and cultured with the same medium (Figure 4A). Remarkably, villus-derived organoids (V-organoids) began forming in only Collagen from day 3 onward (Figure 4B, Figure S2B, C), and 36.4% of them were tdTomato+ (Figure S2D). In contrast, consistent with previous reports^5^, organoid formation was not observed in Matrigel from villus-derived cells (Figure 4B), confirming that the isolated villi did not contain functional CBCs. For comparison, crypt-derived cultures formed organoids within 3 days under both conditions, while villus cultures required a longer time to initiate V-organoid formation (Figure S2C). In primary V-organoids, prominent YAP activation was observed, similar to previously reported Collagen organoids^14, 41^, suggesting that YAP activation may function as an initiating event of fetal reversion in V-organoids (Figure 4C). Once established, V-organoids could be maintained and passaged in both Collagen and Matrigel matrices. The V-organoids raised in Collagen expressed Ly6a, a marker of revSC (Figure 4D) broadly, and upon replating into Matrigel, Olfm4 expression appeared, indicating acquisition of CBC identity (Figure 4E). The resulting budding organoids contained multiple differentiated epithelial lineages including Lyz+ Paneth cells, Muc2+ goblet cells, Dclk1+ tuft cells, Chga+ enteroendocrine cells, and Alpi+ absorptive enterocytes (Figure S2E).

**Figure 4.**
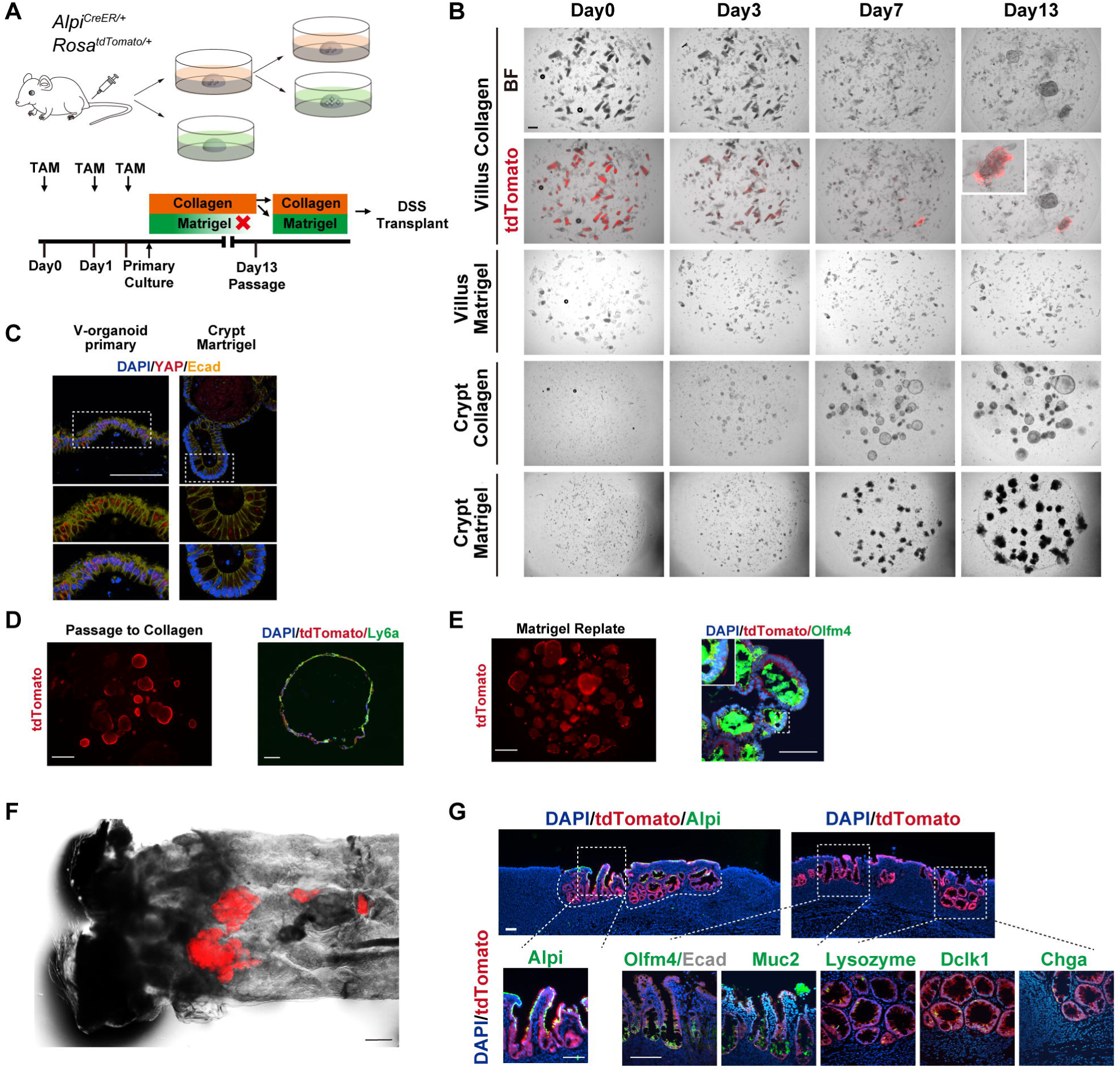
Fetal reversion enables villus-derived cells to regenerate tissue without oncogenic mutations. (A) A schematic presentation of villus culture experiment is shown. Villus and crypt fractions were isolated from the small intestine of *Alpi^CreER/+^; Rosa^tdTomato/+^*, in which Alpi+ enterocytes were labeled by intraperitoneal tamoxifen injection. Isolated villi were cultured in Collagen or Matrigel. V-organoids established in Collagen were subsequently passaged into Collagen or Matrigel and transplanted into DSS-treated colitis model mice. (B) Representative fluorescence images merged with bright field images of villus and crypt cultures in Collagen or Matrigel. In Collagen, tdTomato+ V-organoids, representing progeny of Alpi+ enterocytes were established from villus-derived cells. Scale bar: 500 µm. (C) Representative immunofluorescence images of primary V-organoids and crypt-derived Matrigel organoids (crypts Matrigel) stained with YAP (red) and E-cadherin (Ecad; yellow) antibodies. Prominent nuclear localization of YAP is observed in V-organoids, while nuclear retention of YAP is highly restricted in the budding-domain in crypts Matrigel. Images are counterstained with DAPI (blue). Scale bars:100 µm. (D) Representative fluorescence (left) and immunofluorescence (right) images of tdTomato+ V-organoids passaged into Collagen. V-organoids were successfully maintained in Collagen. Immunostaining revealed high expression of Ly6a (green). Images are counterstained with DAPI (blue). Scale bars: 500 µm (left), 100 µm (right). (E) Representative fluorescence (left) and immunofluorescence (right) images of tdTomato+ V-organoids replated into Matrigel. V-organoids were successfully replated in Matrigel and immunostaining revealed the presence of Olfm4⁺ cells (green). Images are counterstained with DAPI (blue). Scale bars: 500 µm (left), 100 µm (right). (F) A macroscopic image of engrafted patches (red) derived from tdTomato-expressing V-organoids. Scale bar: 1 mm. (G) Immunofluorescence images of cell-specific markers and Alpi staining (green) of serial sections of engrafted patches shown in (E). Transplanted V-organoids regenerated a homeostatic intestinal epithelium while maintaining small intestinal epithelial phenotypes, including Olfm4+ CBCs (green; counterstained with Ecad in gray), Muc2+ goblet cells (green), Lyz+ Paneth cells (green), Chga+ EECs (green), Dclk1+ Tuft cells (green), and Alpi+ absorptive enterocytes (green). Images are counterstained with DAPI (blue). Scale bar: 100 µm.

Finally, to assess the regenerative potential of V-organoids in vivo, we employed a DSS-induced colitis transplantation model^21, 22, 42^. Notably, tdTomato+ V-organoids successfully engrafted in 67% of recipient mice (Figure 4F, Figure S2F). Serial sectioning of engrafted tissue revealed reconstitution of a homeostatic small intestinal epithelium composed of Olfm4+ CBCs, Muc2+ goblet cells, Lyz+ Paneth cells, Dclk1+ tuft cells, Chga+ EECs, and Alpi+ enterocytes (Figure 4G).

Typically, organoids are established from CBC-containing crypts, and villi, which lack stem cells fail to generate organoids in Matrigel. Previous studies have shown that in the presence of oncogenic mutations, villus-derived differentiated cells can acquire stem cell–like properties and form organoids^5, 39, 40^. Notably, our findings demonstrate that, even in the absence of oncogenic mutations, Alpi+ villus enterocytes can undergo fetal reversion to reacquire stemness, enabling their contribution to epithelial regeneration. These findings highlight that the recruitment of diverse epithelial lineages reinforces the robustness of intestinal repair.

### Alpi+ enterocytes replenish the intestinal stem cell pool via fetal reversion in vivo

To further validate whether differentiated enterocytes can undergo fetal reversion and contribute to regeneration in another in vivo setting, we employed a lineage tracing approach in a 5-fluorouracil (5-FU) induced enteritis model^23, 24^. *Alpi^CreER/+^; Rosa^tdTomato/+^* mice were intraperitoneally injected with tamoxifen to label Alpi+ enterocytes, followed by administration of 5-FU to induce intestinal injury (Figure 5A). Histological analysis on day 3 revealed characteristic inflammatory changes, including villus shortening and stromal thickening, as well as marked body weight loss (Figure S3A, B). Notably, Collagen type I (Col I) was prominently upregulated in the stromal compartments of both the crypts and villi (Figure 5B). Correspondingly, the villus compartment, characterized by cytoplasmic retention of YAP, occasionally displayed nuclear localization of YAP, as observed in the crypt domain (Figure 5C). These findings indicate that ECM remodeling and YAP activation—key drivers of fetal reversion^14, 35^—occur concomitantly during the early phase of 5-FU induced enteritis. At this time point, we detected co-expression of Ly6a in tdTomato+ cells localized to the atrophic villi (Figure 5C), suggesting that Alpi+ enterocytes had undergone reprogramming toward a revival stem cell (revSC)-like state^15^. By day 14, tdTomato+ clonal ribbons were observed in the small intestine of 5-FU treated mice, but not in vehicle-treated controls (Figure 5D), supporting the contribution of Alpi+ cells to epithelial regeneration. Quantitative analysis revealed a significantly higher frequency of clonal ribbons in the 5-FU group compared to controls within the first 5 cm intestinal tube from the pylorus (Figure S3C).

**Figure 5.**
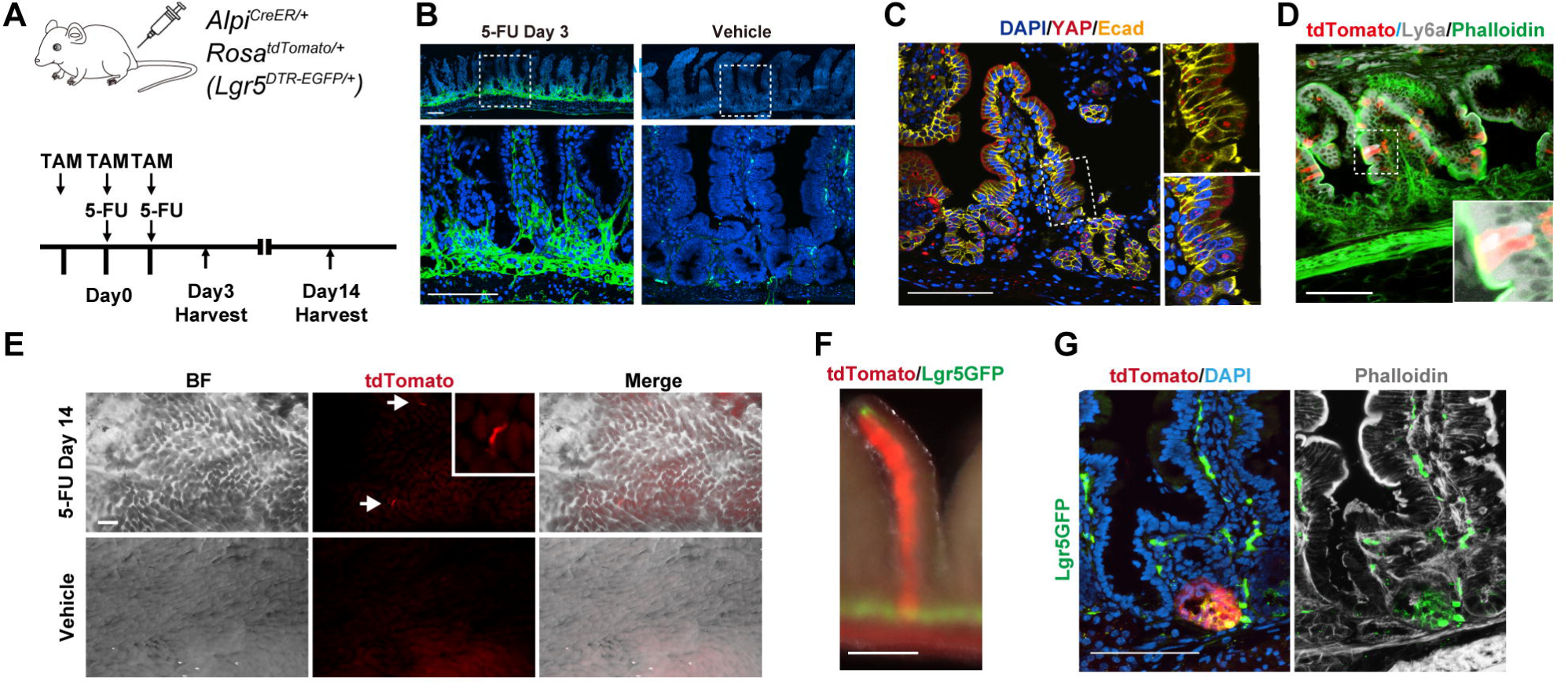
Alpi+ enterocytes replenish the intestinal stem cell pool via fetal reversion in vivo. (A–E) Lineage tracing of Alpi+ enterocytes in a 5-FU–induced chemical enteritis model. (A) Tamoxifen (10 mg/kg) was administered intraperitoneally to *Alpi^CreER/+^; Rosa^tdTomato/+^* mice on days –1, 0, and 1. 5-FU (200 mg/kg) was injected intraperitoneally on days 0 and 1 to induce enteritis. Histological analyses were performed on days 3 and 14. (B) Immunostaining for collagen type I (Collagen I; green) in the small intestine on day 3 after 5-FU treatment or vehicle control. ECM-rich boxed regions are enlarged below. Images are counterstained DAPI (blue). Scale bar: 100 µm. (C) Immunofluorescence staining for YAP (red) in the small intestine on day 3 after 5-FU treatment. Nuclear localization of YAP is observed in both crypt and villus regions. Images are counterstained with Ecad (yellow) and DAPI (blue). Scale bar: 100 µm. (D) Detection of Ly6a expression in the small intestine on day 3 after 5-FU treatment. Boxed regions are enlarged. Co-localization of Ly6a (gray) and tdTomato (red) signals was observed. Images are counterstained with Phalloidin (green). Scale bar: 100 µm. (E) Macroscopic images of the small intestine on day 14 after 5-FU treatment. Brightfield (BF), tdTomato (red), and merged images are shown. Arrows indicate tdTomato+ clonal ribbons, which were observed only in 5-FU–treated mice but not in vehicle-treated controls. Scale bar: 500 µm. (F) A macroscopic image of tdTomato+ clonal ribbons in *Alpi^CreER/+^; Rosa^tdTomato/+^*; *Lgr5^DTR-EGFP/+^* mice on day 14 after 5-FU treatment. TdTomato (red) and GFP signals (green) were co-localized at the base of the crypt. Scale bar: 200 µm. (G) Immunofluorescence images of the tdTomato+ clonal ribbon shown in (E). tdTomato (red) and GFP signals (green) were histologically co-localized at the crypt bottom. Sections were counter stained with Phalloidin (gray) and DAPI (blue). Scale bar: 100 µm.

To further confirm that Alpi+ cells can give rise to CBCs, we utilized *Alpi^CreER/+^; Rosa^tdTomato/+^; Lgr5 ^DTR-EGFP/+^* mice, which enabled simultaneous lineage tracing of Alpi+ enterocytes and visualization of Lgr5+ CBCs to evaluate clonal ribbon formation. Following tamoxifen administration to label Alpi+ cells, mice were subjected to 5-FU induced enteritis as described above. By day 14 observation, co-localization of tdTomato and GFP signals at the crypt base was confirmed both macroscopically and by histological examination (Figure 5F, G), demonstrating that Alpi+ enterocyte-derived cells had reacquired Lgr5 expression and occupied the CBC niche. Collectively, these results indicate that differentiated enterocytes can undergo fetal reversion and regenerate the intestinal stem cell compartment in response to inflammatory injury.

### Fetal reversion renders inflammatory stress tolerant

Given that both CBCs and differentiated cells, including Alpi+ enterocytes, can undergo fetal reversion to become revSCs—a transient and inflammation-associated stem cell state—we hypothesized that revSCs possess a functional advantage in stress tolerance. To test this, we examined the response of organoids to inflammatory stress induced by 5-FU. Mouse and human small intestinal organoids were cultured in either Matrigel or Collagen and treated with 40 µM 5-FU from day 4 to day 6 (Figure 6A). In both species, Matrigel organoids exhibited marked growth inhibition following 5-FU treatment, whereas Collagen organoids retained their proliferative capacity (Figure 6B, 6C).

**Figure 6.**
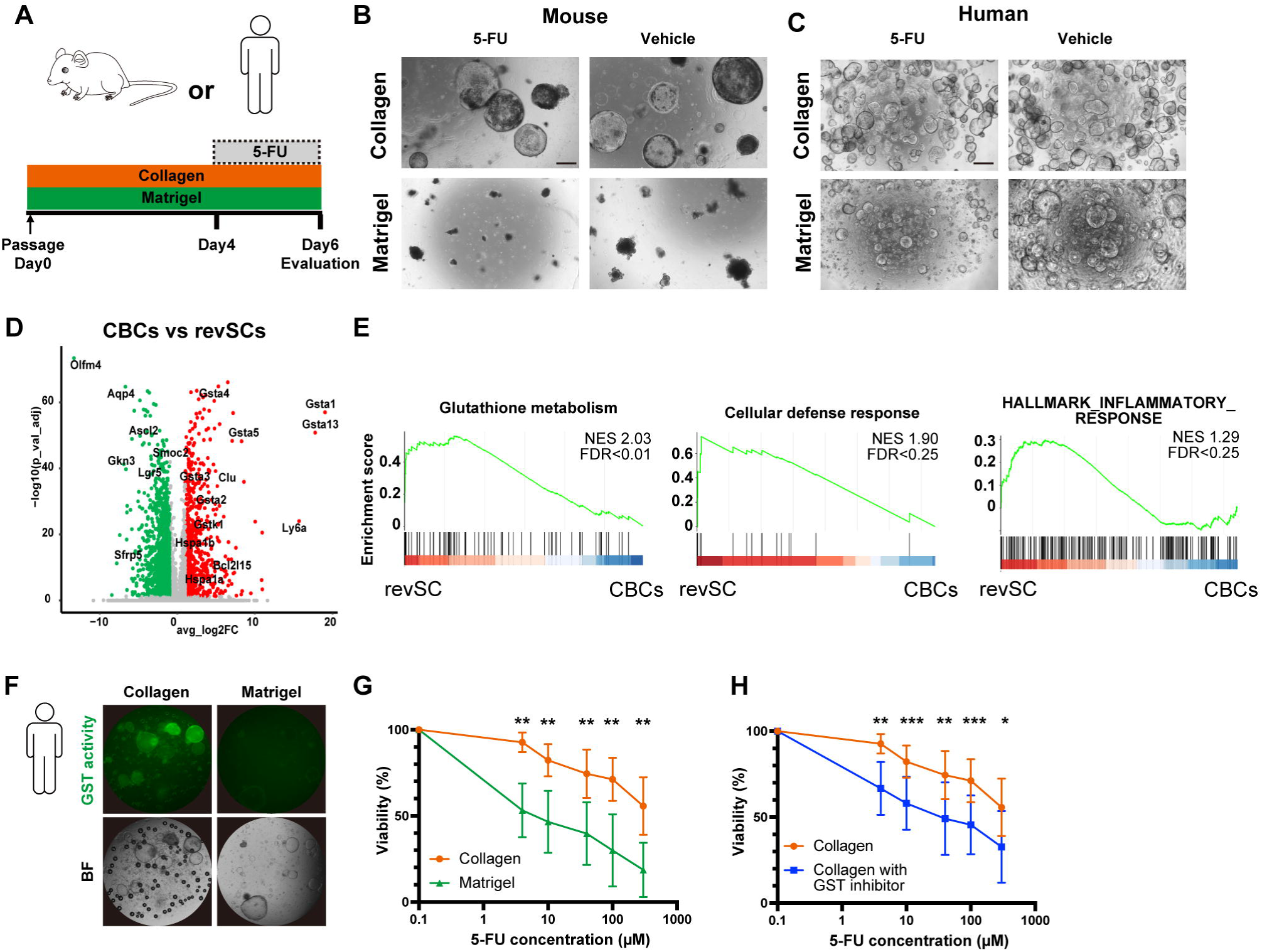
Fetal reversion drives inflammatory stress tolerance. (A) Mouse and human small intestinal organoids were treated with 5-FU (40 µM) from day 4 to day 6 after passage. (B) Representative images of mouse organoids cultured in Matrigel or Collagen matrices on day 6, treated with 5-FU or vehicle. Scale bar: 500 µm. (C) Representative images of human organoids cultured in Matrigel or Collagen matrices on day 6, treated with 5-FU or vehicle. Scale bar: 500 µm. (D) Volcano plot of differentially expressed genes (DEGs; fold change > 1 and \**P* < 0.05) between CBCs cluster and revSCs cluster of mouse organoids. Genes of interests are indicated. (E) Gene set enrichment plots showing significant enrichment of glutathione metabolism (KEGG), cellular defense response (GO:0006968), and HALLMARK inflammatory response (MSigDB) in the revSC gene set of human intestinal organoids. Enrichment analysis was performed using the clusterProfiler package. (F) GST (glutathione S-transferase) activity assay of human organoids cultured in Collagen and Matrigel matrices. GST activity is visualized as green fluorescent signal. Brightfield (BF) images are shown below. (G) Cell viability assay after 48 hours of 5-FU treatment in Collagen and Matrigel cultures. Data are shown as mean ± SD (n = 7, two-tailed Student’s *t*-test, \*\**P* < 0.01). (H) Cell viability assay after 48 hrs of 5-FU treatment in Collagen and Matrigel cultures in the absence or presence of GST inhibitor. Data are shown as mean ± SD (n = 7, two-tailed Student’s *t*-test; \**P* < 0.05, \*\**P* < 0.01, \*\*\**P* < 0.001).

To explore potential mechanisms underlying this resilience, we performed differential gene expression analysis using scRNA-seq data from mouse organoids. Compared to CBCs cluster, revSCs cluster exhibited significant upregulation of genes associated with cellular stress protection, including glutathione S-transferase (GST) family members^43^, Bcl2 family genes^44^, and heat shock protein family genes^45^ (Figure 6D). Gene set enrichment analysis (GSEA) was performed using differentially expressed genes to evaluate pathway-level changes between revSCs and CBCs. Enrichment results highlighted three major pathways: glutathione metabolism (KEGG), cellular defense response (GO), and the HALLMARK inflammatory response signature (MSigDB) (Figure 6E). These pathways represent antioxidant defense, innate immune activation, and stress adaptation, collectively supporting the notion that revSCs are transcriptionally primed for inflammatory resilience.

Given the role of GST in detoxification and therapy resistance, particularly in inflammatory bowel disease (IBD)^46, 47^ and cancer^48, 49^, we directly measured GST activity in human organoids. Collagen organoids exhibited strong GST activity, which was maintained even after 5-FU treatment, whereas Matrigel organoids showed little to no GST activity regardless of treatment (Figure 6F). Cell viability assays further demonstrated that Collagen organoids were significantly more tolerant to 5-FU than their Matrigel counterparts (Figure 6G). Notably, inhibition of GST activity in Collagen organoids restored sensitivity to 5-FU, confirming the functional relevance of glutathione metabolism in mediating stress resilience (Figure 6H). These findings highlight the critical role of fetal reversion in endowing intestinal epithelial cells with enhanced tolerance to inflammatory insults in a GST-dependent manner.

## DISCUSSION

In this study, we investigated the interplay between spatial plasticity and fetal reversion in the intestinal epithelium, revealing a stress-adaptive regenerative hierarchy in which revSCs emerge from diverse origins—including CBCs and enterocytes—to preserve stem cell function under injury (Figure 7). While the intestinal epithelium regenerates via both spatial plasticity of differentiated cells and fetal reversion during inflammation, how these modes interact remains unclear. In particular, the hierarchical relationship among differentiated cells, CBCs, and revSCs, and the biological significance of their transitions, has yet to be fully understood.

**Figure 7.**
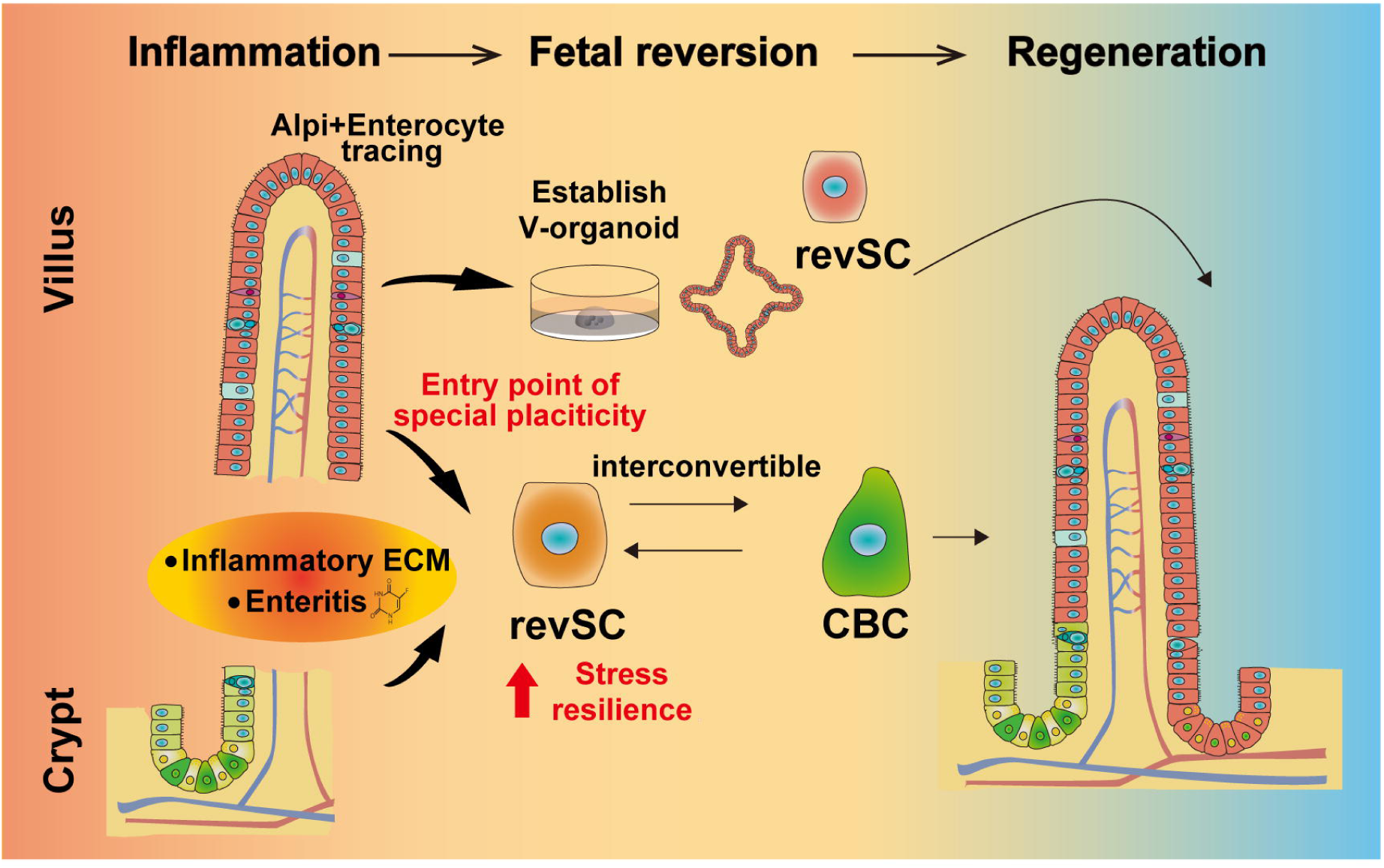
Schematic illustration of the role of revSCs in intestinal epithelial regeneration. In response to inflammation, both villus-derived differentiated enterocytes and crypt-derived CBCs convert into revSCs and contribute to epithelial regeneration. Fetal reversion enables the formation of villus-derived organoids (V-organoids) without oncogenic mutations. revSCs can also revert to CBCs and regenerate the entire intestinal epithelium. Our findings establish fetal reversion as a cellular escape mechanism that safeguards epithelial regeneration under inflammatory conditions.

To address these questions, we systematically analyzed the cellular origins and functional roles of revSCs. First, our results suggest that the majority of revSCs are supplied by CBCs. Previously, the transient nature of revSCs has posed a barrier to their functional characterization, with current understanding largely based on in silico estimations derived from single-cell transcriptomic analyses^31,35^. In this study, by establishing and utilizing a revSC-rich organoid culture system, we successfully provided experimental evidence demonstrating the high fetal reversion potential of CBCs. This finding reinforces the central role of homeostatic stem cells in maintaining intestinal integrity, even under regenerative conditions. Also, we showed that revSCs possess robust stemness and the ability to give rise to CBCs across various models: organoid cultures, organoid transplantation^22, 42^ and the 5-FU enteritis model. Importantly, the CBCs derived from revSCs were functionally indistinguishable from endogenous CBCs under homeostatic conditions, supporting the notion that revSCs act as bona fide stem cells capable of restoring the homeostatic stem cell pool.

Moreover, we demonstrated that revSCs could also arise from Alpi+ differentiated enterocytes, both in vitro and in vivo, indicating that multiple cellular sources contribute to the revSC pool. This finding extends previous reports identifying Clusterin+ cells and p57+ entecoendocrine cells as potential origins of revSCs, by revealing differentiated enterocytes— the most abundant cell population in the intestinal epithelium—as an additional and functionally significant source. Given the scarcity of the previously proposed revSC-originating populations, the involvement of enterocytes may represent a robust regenerative mechanism that ensures reliable epithelial repair under inflammatory conditions.

Furthermore, we revealed that villus-derived cells can form organoids via fetal reversion when cultured in Collagen matrix, without oncogenic mutations. This finding challenges the long-standing dogma that intestinal organoids can only be established from crypts containing stem cells^25, 26^, and that villus-derived cells lacking stem cells are incapable of organoid formation. To date, the only reported exception has involved cells harboring oncogenic mutations, which can acquire stem cell–like properties and give rise to organoids from villus epithelium^39, 40^. In contrast, our study demonstrates that fetal reversion enables differentiated villus cells to regain stemness through mechanisms independent of genomic mutations. Importantly, our observation that tdTomato-negative organoids can also emerge from villus cultures supports the notion that multiple differentiated epithelial cell types beyond the Alpi+ lineage may contribute to revSC generation. While our current analysis focused specifically on differentiated enterocytes, further investigations are required to elucidate the full spectrum of contributing differentiated populations and to define the hierarchical relationships that govern their regenerative potential. Our V-organoid system may serve as a powerful platform to further explore how various differentiated cell types undergo fetal reversion and contribute to regenerative responses under inflammatory or injury conditions.

Additionally, we found that revSCs exhibit high GST activity and resistance to 5-FU treatment, reflecting a high degree of stress tolerance. The emergence of revSCs during inflammation thus may represent a protective mechanism that enables stem cells to withstand hostile environments. These novel insights into a stress-adaptive regenerative framework not only redefine intestinal epithelial repair, but also open new avenues for understanding inflammation-associated diseases—such as IBD and colorectal cancer (CRC)—by optimizing treatment strategies and exploring fetal-like regenerative responses across organ systems. Importantly, our findings offer critical insights into fetal reversion that may inform the understanding and treatment of inflammation-associated pathologies. Recent single-cell transcriptomic studies have identified revSC-like populations as inflammation-specific epithelial subsets, highlighting their significance in tissue regeneration^15, 16^. In light of our findings, the maintenance of epithelial integrity via revSCs may represent a rational escape strategy under inflammatory stress. Moreover, the selective emergence of stress-tolerant revSCs under sustained inflammation may reflect an adaptive response to a hostile tissue environment. Conversely, there is growing evidence that failure to properly resolve fetal reversion may contribute to inflammation-associated tumorigenesis^50^. Therefore, elucidating the behavior of revSCs in human disease contexts will be critical for improving IBD and inflammation-associated tumorigenesis treatment strategies.

Furthermore, fetal reversion, particularly in the context of therapy resistance and CRC progression, has come to be recognized as a form of “onco-fetal reprogramming”^17, 18^. We previously reported that ECM-dependent fetal reversion contributes to chemoresistance in CRC cells^51^. Non-malignant revSCs also exhibit 5-FU resistance, indicating that fetal reversion–associated traits are not solely mutation-driven. This distinction may help to clarify the mechanisms of therapy resistance and to guide the development of novel therapeutic strategies in CRC. While further validation in human disease contexts remains necessary, our scRNA-seq data from human Collagen organoids suggest that this system holds promise for translational investigation.

The traits of revSCs—namely, their flexible supply from diverse cellular origins and their resilience to stress—highlight their potential as a therapeutic target in regenerative medicine. Similarly, TGF-β–induced fetal reversion has been shown to enhance epithelial repair in transplantation models^31^, further exemplifying the therapeutic relevance of fetal-like reprogramming. Recent studies have also shown that revSCs can also be induced by multiple factors such as TGF-β^31^, Asporin^52^ and IFN-γ^13^. As exemplified by our Collagen organoid model, analyzing individual factors in isolation can help simplify and deepen our understanding of such complex regenerative mechanisms. Also, in physiological settings, fetal reversion is likely orchestrated by a combination of multiple signals that together induce dynamic changes in cell states. Future investigations should now focus on how these pathways operate within a unified, systems-level framework, which may ultimately enable the development of more precisely targeted and optimized therapeutic strategies. Notably, recent studies have described fetal-like reprogramming—termed “paligenosis”—as a conserved regenerative response in multiple organs^53, 54, 55, 56^. These parallels suggest that transient reversion to a developmental state may represent a general strategy for promoting tissue repair under stress, offering a potential universal target for regenerative therapies.

## MATERIALS AND METHODS

### Mice

*C57BL/6J* wild type mice were obtained from CLEA Japan (Tokyo, Japan). *Lgr5-^EGFP-IRES-CreERT2^* (Stock#:008875)^2^ mice and *CAG-LSL-tdTomato* mice (Stock#:007914)^57^ were obtained from The Jackson Laboratory. *Lgr5 ^DTR-EGFP^* ^37^ mice were kindly provided by F. de Sauvage. *ALPi^CreER^* ^5^mice were kindly gifted by Hans Clevers. All animal studies were approved by the Institutional Animal Care and Use Committee of Institute of Science Tokyo (No. A2025-020C).

### Mouse treatment by Tamoxifen and Fluorouracil (5-FU)

To trace ALP+ enterocytes in vivo, *Alpi^CreER/+^; Rosa^tdTomato/+^* mice were administered with Tamoxifen (10mg/kg body weight; Sigma-Aldrich # H6278) twice on day 0 and day 1 by intraperitoneal administration. On day 2, small intestinal crypts were harvested for primary culture either in Matrigel and collagen type I as described below. *Alpi^CreER/+^; Rosa^tdTomato/+^*; with or without *Lgr5^DTR-EGFP/+^* mice were also treated with Tamoxifen three times on day −1, day0 and day 1 for continuous 3 day in order to fully label ALP+ enterocytes before induction of enteritis, which was initiated by intraperitoneal administration of 5-FU;(200mg/kg body weight; Sigma-Aldrich, #F6627) twice on day 0 and day 1.

### Organoid culture of adult murine small intestinal epithelium in Matrigel or Collagen (For mainly Figure 1,3,6)

Mouse small intestinal organoids were established and maintained as previously described^14^. Briefly, small intestines of C57/BL6 mice or *Alpi^CreER/+^; Rosa^tdTomato/+^* mice were harvested from adult mice, flushed with cold phosphate buffer saline (PBS; Nacalai Tesque; #14249-24), and opened longitudinally. Tissues were cut into 2–5 mm fragments and incubated in 2 mM ethylenediaminetetraacetic acid (EDTA; Invitrogen; #AM9260G) in PBS at 4 °C for 30 minutes to dissociate the epithelial layer in a 50 mL conical tube (VIOLAMO, #1-3500-22). The epithelial fractions were collected by vigorous shaking, filtered through a 70 μm cell strainer (Falcon; #352350), and centrifuged at 500 × g for 3 minutes after transfer into a 15 ml conical tube (Falcon, #188271-013). After washing, the resulting crypts were resuspended in either Matrigel (Corning, #356231) or collagen type I (Cellmatrix Type I-A; Nitta Gelatin, #631-00651) and plated as 30 μL domes in pre-warmed 24-well plates (Corning, #3524).

After polymerization at 37 °C for 30 minutes, domes were overlaid with 500μL of ENR medium, consisting of Advanced DMEM/F12 (Gibco, #12634-010) supplemented with Penicillin/Streptomycin (1% v/v; Nacalai Tesque; #26253-84), GlutaMAX (1% v/v; Gibco, #35050-061), 1X B27 supplement (Gibco, #17504-044), 1X N2 supplement (Gibco, #17502-048), recombinant murine EGF (50 ng/mL; PeproTech, #315-09), recombinant murine Noggin (100 ng/mL; PeproTech, #1967-NG), and recombinant murine R-spondin1 (400 ng/mL; Qkine, QK006). For Collagen culture (Figure1), murine Wnt3a (100 ng/ml; Abcam, #ab81484), Nicotinamide (10 mM; Sigma-Aldrich, #72340), and bovine serum albumin (1% v/v; BSA; Sigma-Aldrich, # A9576) were supplemented to ENR medium (ENRW). For initial seeding, Y-27632 (10 µM; Tocris, #1254) was added for the first 2-3 days. Medium was changed every 2–3 days.

Occasionally, for single cell culture (Figure 2), replating to Matrigel (Figure 2,3), and villus culture (Figure 4), comparison in 5-FU treatment (Figure 6), 2**5**% cWRNE consisting of Advanced DMEM/F12 supplemented with Penicillin/Streptomycin (1% v/v), GlutaMAX (1% v/v), 25% (v/v) L-WRN conditioned medium ^58^ concentrated using 10 kDa Amicon Ultra filter unit (Merck, #UFC901008), murine EGF (50 ng/mL), 1X N2 supplement, 1X B27 supplement, N-Acetyl-L cysteine (1 mM; Sigma-Aldrich, #A9165) were used.

### Primary culture of human proximal small intestinal epithelial cells in Matrigel or Collagen type I (For Figure 1 and 6)

Human proximal small intestinal crypts were harvested from surgical specimens. The epithelium was dissected and cut into small pieces. The fragments were washed in 30ml PBS twice and subsequently in 2% Mucofilin (Eisai) once, followed by 4°C incubations in 15 mM EDTA for 20 minutes. After washing in 30ml PBS, the tissues were incubated in 10ml collagenase solution (Sigma-Aldrich; 0.5 mg/mL in PBS, #C7657) at 37°C for 20 minutes. After settling down, the supernatant was transferred into another 50 mL conical tube by passing through a 70 µm cell strainer. The total volume was adjusted to 10 mL with 0.1%(v/v) BSA/PBS into 15 mL conical tube and cells were pelleted at 500 × g for 3 min. The pellets were suspended in Matrigel and plated onto a 24-well plate. Organoids were cultured in 20% cWRNE medium consisting of Advanced DMEM/F12, supplemented with Penicillin/Streptomycin (1% v/v), GlutaMAX (1% v/v), 20% (v/v) L-WRN conditioned medium, murine EGF (50 ng/mL), Nicotinamide (10 mM), 1X N2 supplement, 1X B27 supplement, N-Acetyl-L cysteine (1 mM), Gastrin I (10 nM; Sigma-Aldrich, #G9145) and A83-01 (500 nM; Tocris, #2939). We passaged Matrigel organoids into Collagen matrix. Medium for Collagen culture use the same above. Surgical specimens were obtained at Institute of Science Tokyo Hospital. The Scientific Ethics Committee of Institute of Science Tokyo Hospital approved the use of this material for research purposes (M2023-195), and informed consent was obtained from all patients.

### Flowcytometory and Cell Sorting for reconstitution assay in *Lgr5^EGFP-IRES-CreERT2/+^ mice* and *Lgr5^DTR-EGFP/+^* organoids (For Figure 2)

Flow cytometric analysis and fluorescence-activated cell sorting (FACS) were performed using BD FACSAria III cell sorter (BD Biosciences) or FACS Melody (BD Biosciences) fitted with a 100 μm nozzle, and the data were analyzed by FlowJo_V10 software. For sorting or analysis of live cells from mouse small intestine, intestines were flushed with cold PBS, opened longitudinally, and cut into 2–5 mm fragments. The tissue was incubated in 2 mM EDTA in PBS for 30 minutes at 4°C with gentle agitation to release the epithelial layer.

The detached epithelial cells were collected by vigorous shaking, filtered through a 70 μm cell strainer. Both these isolated crypts, and mechanically dissociated organoids, were enzymatically dissociated into single cells using TrypLE Express (Gibco; #12605-010) at 37°C for 20 minutes with intermittent pipetting.

Epithelial cells were gated as EpCAM+ (eBioscience, #17-5791-80), CD45- (Biolegend, #103133), CD31- (Biolegend, #102423), Ter119- (Biolegend, #116233) and 7AAD- (Biolegend, #420403). Lgr5-GFP fluorescence intensity was used to classify cells into GFPhigh, GFPlow and GFPneg populations as previously reported^25^. Epithelial cells from *Lgr5^DTR-EGFP/+^* organoids were gated as EpCAM⁺CD45⁻CD31⁻Ter119⁻7AAD⁻, and Lgr5-EGFP fluorescence intensity was used to classify cells into GFPhigh and GFPneg populations, where the GFPneg gate was defined based on the background fluorescence intensity of WT control organoids lacking EGFP expression. For single-cell dissociation from organoids, Matrigel was removed using Cell Recovery Solution (Corning, #354253), while collagen-based matrices were digested using collagenase solution (0.625 mL in PBS). The collected organoids were then incubated in TrypLE Express for 15–30 minutes. Mechanical dissociation was performed using pipetting or a 5 mL syringe to obtain single-cell suspensions. Finally, the dissociated cells were filtered through a 40 µm cell strainer (Falcon, #352340) to remove debris and aggregates. Ly6a antibody (Biolegend, #108111) was used for the evaluation of Ly6a expression. Sorted populations were collected into cooled Advanced DMEM/F12 and 5000 live cells were suspended in either Matrigel or Collagen matrices and plated as 20 μL domes in pre-warmed 24-well plates. In need, glass bottom 24-well culture plate was used for better imaging of organoids (Iwaki, #637-35161). For following culture, 100 ng/mL of murine Wnt3a, 100 ng/ml of murine recombinant Noggin, 400 ng/mL of murine recombinant R-spondin1, 50 ng/mL of murine EGF and 1 µM of Jagged-1 (AnaSpec; # 188-204) were mixed in either Matrigel or Collagen matrices, and 25% cWRNE medium described in the previous section with additional 3 µM of Chir99021(Stemgent, # 04-0004-10) and 2.5 µM of Thiazovidin (Stemgent, #04-0017) was used (reconstitution medium).

### Villus spheroid culture from *ALPi^CreER/+^; Rosa^tdTomato/+^* (For **Figure 4**)

*Alpi^CreER/+^; Rosa^tdTomato/+^* mice were intraperitoneally injected with tamoxifen (10 mg/kg body weight) on days 0, 1, and 2 (12 hours before isolation) to label Alpi+ lineages. Villus isolation was performed as previously described ^5, 39, 40, 59^. Briefly, small intestines were flushed with PBS and opened longitudinally. Villi within 3 cm distal to the pylorus were scraped using a glass coverslip (MATSUNAMI, #MAS-01), washed in PBS, and centrifuged at 100 × g for 3 minutes to separate villi from single cells. The collected villi were further washed using a 70 µm cell strainer placed into one well of a 6-well plate (Falcon, #353046) filled with PBS and gently pipetted. The strainer was then transferred sequentially into other wells filled with fresh PBS to ensure thorough washing. After washing, PBS and villi remaining within the strainer were collected into a 15 mL tube and centrifuged at 500 × g for 3 minutes. Collected villi were suspended in 20 µl of either Collagen or Matrigel matrices and plated in 24-well plates. Cultures were maintained in 25% cWRNE medium. V-organoids established in Collagen were passaged into either Collagen or Matrigel matrices, using the same method as crypt-derived organoids, and cultured in cWRNE or ENR medium. To isolate tdTomato-labeled V-organoids, tdTomato+ organoids were manually identified under a fluorescence stereomicroscope, and surrounding Collagen matrix was excised using a razor blade (Feather. #FAS-10).

### Organoids treatment by diphtheria toxin (DT), 4-hydroxy Tamoxifen (4-OHT)

Small intestinal organoids derived from *Lgr5^DTR-EGFP/+^* mice were treated by DT (0.05 ug/mL; Sigma #D0564) in vitro to delete Lgr5+ CBC before passaging to the next generation. Small intestinal organoids derived from *Alpi^CreER/+^; Rosa^tdTomato/+^* mice were treated by 4-hydroxytamoxifen (4-OHT; 100 µM; Sigma-Aldrich, #SML1666) to label and trace alp+ enterocytes in vitro.

### 5-FU inflammation model in vitro

Human or mouse Matrigel organoids or Collagen organoids were seeded on 48-well plates (Thermo Fisher Scientific, #150687) in cWRNE medium. From Day 4-6, 5-FU (4 µM-300 µM; Selleck Chemicals, #S-1209) was added to the culture medium at various concentrations to compare the resilience for inflammatory stress between Matrigel organoids and Collagen organoids. For GST inhibition, on Day 4, immediately prior to the initiation of 5-FU treatment, the GST inhibitor (Funakoshi, #FDV0031) was diluted in HBSS+ (Nacalai Tesque, # 17459-55) to a final concentration of 10 µM and incubated at 37 °C for 60 minutes. Following incubation, the inhibitor was added to the 5-FU–containing culture medium to reach a final concentration of 100 µM. On day 6, organoids were subjected to cell viability and GST activity assays.

### Cell Viability Assay

Cell viability was assessed at day 6 of 5-FU inflammation model using the CellTiter 96® AQueous One Solution Cell Proliferation Assay (Promega; # G3582) according to the manufacturer’s instructions. Absorbance was measured at 490 nm using a GloMax® Discover Microplate Reader (Promega; # GM3000).

### GST activity assay

GST activity was measured using the CellFluor GST (Funakoshi; #FDV-0030) following the manufacturer’s instructions. Organoids cultured in 96-well plates (Falcon, #353072) were washed once with PBS, and the culture medium was replaced with a 10 µM working solution prepared in HBSS+. The plates were incubated at 37 °C for 1 hour. After incubation, organoids were washed again with PBS and imaged using a Keyence BZ-X810 fluorescence microscope to detect green fluorescence signals.

### Bright field and fluorescence images of in vitro organoids

Bright field and fluorescence images acquired using a Olympus CKX53 equipped with Olympus DP74 CCD camera and U-LGPS Fluorescence Light Source or Keyence BZX810.

### Organoids transplantation

Transplantation was performed using C57BL6 mice as recipients, essentially as previously described^21, 22, 42^. Briefly, Dextran Sulfate Sodium (DSS; MP biomedicals, #160110) was administered to *C57BL6* mice (3 to 6 months old) for 5 days. Animals were infused with cultured intestinal epithelial cells derived from villus derived organoid of *Alpi^CreER/+^; Rosa^tdTomato/+^* mice on day 8 after the beginning of DSS treatment. V-organoids were cultured as described above in Matrigel after establishment in Collagen. Epithelial cells were released from matrix and mechanically dissociated into sheets of epithelial cells. After washing, cell fragments from approximately 1000 organoids were resuspended in 300μL of 5% Matrigel in PBS. A flexible catheter (TERUMO, #SV-21CLK) was inserted into the colon of mice under general anesthesia (Isoflurane; VERITAS), and the cell suspension was subsequently infused into the colonic lumen. The anus was sealed with surgical histoacryl glue (B. Brawn), which was removed after 3 hours. After transplantation, animals were carefully monitored during recovery. Two weeks after the transplantation, recipient mice were sacrificed, and colons were harvested for analyses. Images of engrafted patch were acquired using a fluorescence stereomicroscope (Olympus, MVX10) in the isolated colon tissues.

### Histological analysis

Tissues were harvested at the indicated time points and fixed in 4% paraformaldehyde (PFA; Nacalai Tesque, #09154-85) in PBS at 4°C overnight. Fixed tissues were incubated overnight at 4°C in 20% sucrose in PBS for cryoprotection, embedded in OCT compound (SAKURA, # SCS-N22), and snap-frozen on liquid nitrogen. Cryosections were cut at a thickness of 8 µm using a cryostat. For permeabilization, sections were incubated in 0.5% Triton X-100 (Nacalai Tesque, #35501-02) in PBS for 10 minutes at room temperature. Sections were incubated in blocking buffer (Nacalai Tesque, #12967-95) consisting of 0.1% Triton X-100 in PBS for 1 hour at room temperature to prevent nonspecific antibody binding. Primary antibodies were diluted in PBS containing 5% blocking buffer and applied to the sections overnight at 4°C. The following primary antibodies were used: anti-Sca-1 (Ly6a) (1:500, R&D Systems, #MAB1226), anti-Olfm4 (1:1000, Cell Signaling Technology, #39141), anti-Lysozyme (1:500, Dako, #A0099), anti-tdTomato (1:200, Origene, #AB8181), anti-Dclk1(1:100, Abgent, #AP7219B), anti-Chga(1:1000, Diasorin, #20085), anti-Collagen I (1:200, Abcam, #ab270993) and anti-YAP(1:200, Cell signaling, #4912). Following primary antibody incubation, sections were washed three times in PBS and incubated with species-specific secondary antibodies conjugated to Alexa Fluor dyes (1:500; Thermo Fisher Scientific) for 1 hour at room temperature in the dark. Nuclei were counterstained and mounted using Fluoro-KEEPER Antifade Reagent (Nacalai Tesque, Cat# 12745-74). Images were acquired using SP8 (Leica) or FV3000 (Olympus) confocal microscope, and were analyzed in Fiji^60^ and Adobe Photoshop CC.

### Alpi staining

To visualize endogenous Alpi activity, enzymatic histochemistry was performed using the ImPACT® Vector Red alkaline phosphatase substrate (Vector Laboratories, SK-5105). Images were acquired using Keyence BZ-X810 microscope, and were analyzed in Adobe Photoshop CC. Upon need, nuclear were counterstained by Hematoxylin (Vector Laboratories, #H3404).

### Hematoxylin and Eosin (H.E.) staining

The rolled intestine of 5-FU enteritis and control tissue was subjected to H.E. staining (Vector Laboratories, #H3502). The images were captured by Keyence BZ-X810 microscope.

### scRNA-seq analysis and data processing

Single-cell suspensions of organoids were prepared according to the protocol described above. For mouse samples, cells were incubated with biotinylated anti-EpCAM antibody (Biolegend, #118203); for human samples, biotinylated anti-CD298 (Miltenyi Biotec, #130-101-292), and anti-CD147(Miltenyi Biotec, # 130-107-101), antibodies were used as primary antibodies. To enable multiplexed single-cell analysis, TotalSeq-A hashtag conjugated to PE-Streptavidin were applied: A0951 (Biolegend, #405251), A0952 (Biolegend, #405253), or A0953 (Biolegend, #405255), allowing for the distinction of experimental conditions. Single-cell suspensions of mouse and human intestinal organoids were processed using the BD Rhapsody system (BD Biosciences) with the BD Rhapsody Targeted & AbSeq Reagent Kit (BD Biosciences, # 633771), according to the manufacturer’s protocol. Captured transcripts were reverse-transcribed on-bead and treated with Exonuclease I for 60 minutes at 37°C. The resulting beads were then subjected to TAS-Seq workflow ^61^ for whole-transcriptome and hashtag cDNA amplification. Briefly, beads were incubated with terminal deoxynucleotidyl transferase (TdT; Roche, # 3333574001) and RNase H (Enzymatics, # Y9220F), followed by second-strand synthesis and the first round of whole-transcriptome amplification (WTA). cDNA was purified using 0.65× AMPure XP beads (Beckman Coulter, # A63882), and hashtag cDNA was isolated from the unbound fraction by adding an additional 0.7× AMPure XP beads (final 1.35×). Second-round amplifications were performed separately for the cDNA and hashtag libraries, followed by further purification with AMPure XP beads. Amplified cDNA libraries were prepared using the NEBNext Ultra II FS DNA Library Prep Kit for Illumina (New England Biolabs, # E7805), and sequenced on an Illumina NovaSeq 6000 system (Illumina, # 20012850) with either NovaSeq 6000 S4 Reagent Kit v1.0 (Illumina, # 20027466) or v1.5 (Illumina, # 20028313), generating 200-cycle paired-end reads (read1: 67 bp, read2: 155 bp). Paired-end Fastq files (R1: cell barcode reads, R2: RNA reads) were processed with Cutadapt (v4.1) for adapter trimming, quality filtering, and phase-shift base removal. Filtered reads were mapped to the mouse or human reference genome (GRCm39 or GRCh38, Ensembl release 107) using STARsolo (v2.7.10a) with BD Rhapsody-specific parameters. Count matrices were generated using STARsolo and DropletUtils (v1.18.1) for knee-plot thresholding. Background droplets were further filtered using Dropkick ^62^. Hashtag assignment and doublet exclusion were performed using flowDensity and custom R scripts. DBEC correction was applied using FlowTrans and mclust packages as described previously. Single-cell RNA-seq analysis was performed using R (v4.3.2) with the Seurat package (v5.1.0)^63^. Raw count matrices were processed via the rDBEC package^61^ to remove background noise and demultiplex BD Rhapsody-derived WTA data. As quality control, doublets were filtered out. Data were log-normalized (NormalizeData, scale.factor = 1,000,000), and variable genes were identified using FindVariableFeatures. To mitigate the effect of cell cycle heterogeneity and the difference in read counts, cell cycle scores were calculated using CellCycleScoring function. Read counts and the difference between the G2M and S phase scores were regressed out by the ScaleData function. Principal component analysis was performed, and significant principal components (PCs) were selected based on the JackStraw method (*p* < 0.05). Clustering was performed using FindClusters (resolution = 0.5 for mouse and 0.3 for human), and UMAP was used for visualization. Marker gene expression and cell type annotation were visualized using FeaturePlot, and DotPlot. Module scores were calculated via AddModuleScore. RNA velocity analysis was performed with velocyto.R (v0.6)^64^ and Python scVelo (v0.2.5)^30^ via the reticulate package (v1.40.0). Differentially expressed genes were defined as those whose *p* value, as calculated by the Wilcoxon rank sum test and adjusted by the Bonferroni method is <0.05 and whose log2[Fold change] is >1 or <−2. GSEA were conducted by utilizing the R software package clusterProfiler (v4.10.1) ^65^ Gene sets from KEGG, Gene Ontology (GO), and the MSigDB Hallmark collection were used as reference databases.

### Statistical analysis

Statistical analyses were performed using GraphPad Prism version 10 (GraphPad Software, San Diego, CA). Data are presented as mean ± SD unless otherwise indicated. For comparisons among three or more groups, one-way ANOVA followed by Tukey’s multiple comparisons test was used. For two-group comparisons, two-tailed Student’s *t*-test were applied. *P*-values were interpreted as follows: \**P* < 0.05, \*\**P* < 0.01, \*\*\**P* < 0.001, \*\*\*\**P* < 0.0001; *P* ≥ 0.05 was considered not significant (n.s.).

## Supporting information

supplementary infomations

## Data availability

The single-cell RNA sequencing data generated in this study related to this manuscript will be deposited upon publication.

## Materials availability

Human organoids used in this study can be used only in Institute of Science Tokyo due to its ethical regulation.

## Acknowledgements

We thank all members of the Department of Gastroenterology and Hepatology, Institute of Science Tokyo for their feedback on this study. We thank Professor Ichiro Sekiya, Dr. Hisako Katano, and all other members of the Center for Stem Cell and Regenerative Medicine, Institute of Science Tokyo for their feedback on this study. We thank Professor Keiichi Nakayama and Dr. Tsunaki Higa at Anticancer Strategies Laboratory, Advanced Research Initiative, Institute of Science Tokyo for their feedback on this study. We thank F. de Sauvage (Genentech Inc.) for providing Lgr5^DTR-EGFP^ mice. FACS sort analysis and preparation of scRNA seq library by BD rhapsody were performed at the Research Core of Institute of Science Tokyo. This research was supported by MEXT/JSPS KAKENHI (23H02887 to S. Yui, 22H00472 to M. Watanabe.), Japan Science and Technology Agency (JST) Forrest Program (JPMJFR2012 to S. Yui), Naoki Tsuchida Research Grant to S. Yui, JST SPRING (JPMJSP2120 and JPMJSP2180 to S. Kirino).

## Author contributions

Conceptualization, S. Kirino., S.Y.; Methodology, S. Kirino, S. Kobayashi, N.O. S.W., F.U., R.O., S.Y.; Investigation, S.Kirino., N.O., S.W., F.U., H.Y., G.I., S.Y.; Analysis, S.Kirino., K.M., Resources, S. Kirino, F.U., S.Y., K.A., D.B., J.H.v.E., C.H.; Writing the draft, S. Kirino., S.Y.; Supervision, M.W., R.O.; Funding; S.Kirino., M.W., S.Y.

