## supplementary infomations for "Fetal reversion from diverse lineages sustains the intestinal stem cell pool and confers stress resilience"

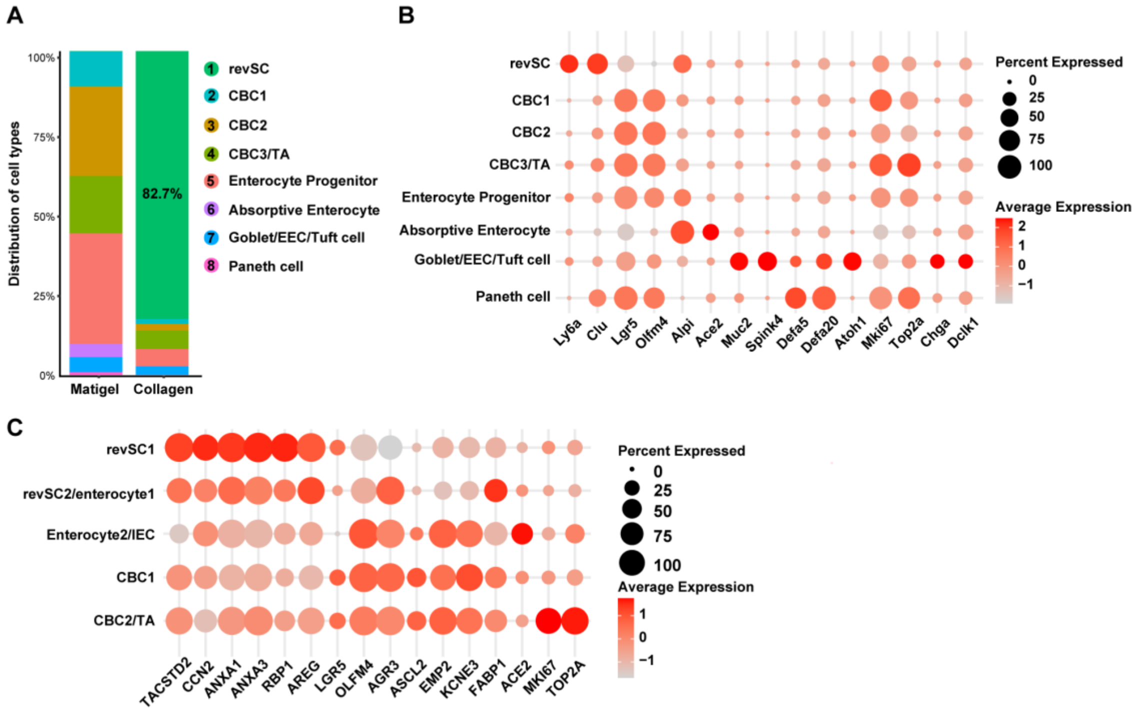


**Figure S1: Cluster details of scRNA seq analysis. Related to Figure 1.**

**(**A) Bar plot showing the distribution of annotated cell types across samples.

(B) Dot plot visualizing the expression of cell type–specific marker genes across clusters in scRNA seq analysis in mouse organoids.

(C) Dot plot visualizing the expression of cell type–specific marker genes across clusters in scRNA seq analysis in human organoids.

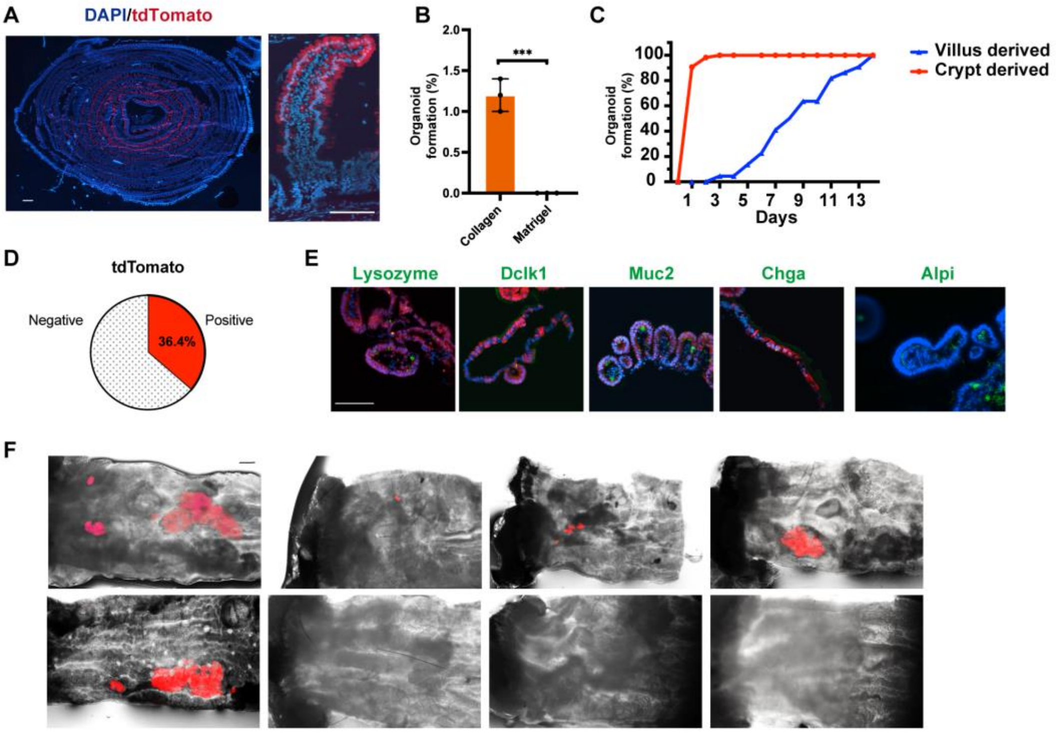


**Figure S2: Details of labelling of ALPi+ enterocytes and villus culture. Related to Figure 4.**

(A) Representative images of the small intestine from *Alpi*$^{CreERKI/+}$; *Rosa*$^{tdTomato\ KI/+}$ mice following tamoxifen induction. The tissue was sectioned in a Swiss-roll configuration, with the proximal region oriented at the center and the distal region at the periphery. Low-magnification image (left) and high-magnification image (right) are shown. Scale bars: 1 mm (left), 100 μm (right).

(B) Quantitative comparison of organoid formation rates from villi cultured in COL and MG. Data are presented as mean ± SD (n=3 per group, two-tailed Student's *t*-test, ***$P$ < 0.001.)

(C) Cumulative graph showing the progression of organoid formation over time. Data were normalized such that the final value on day 13 was set to 100%.

(D) Percentage of tdTomato+ organoids among V-organoids (n = 11 V-organoids).

(E) Immunofluorescence and Alpi staining images of the V-organoid replated in MG shown in Figure 4D, which is entirely composed of tdTomato+ cells. Scale bar: 100 μm.

(F) Macroscopic images of engrafted patches derived from villus-derived organoids, distinct from those shown in Figure 4E.

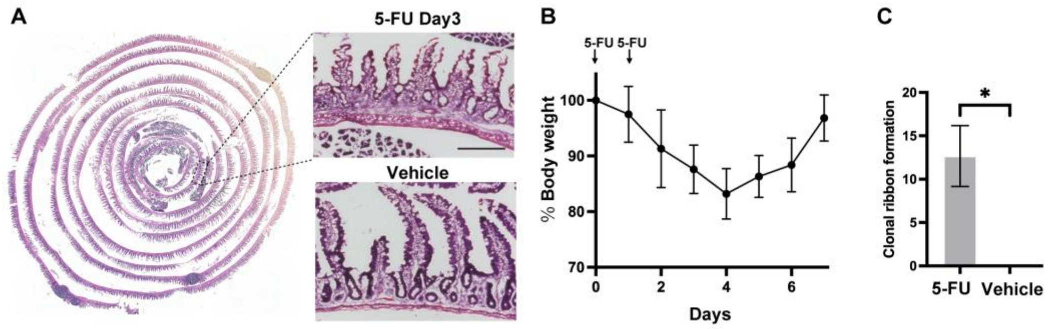


**Figure S3: Supportive information of 5-FU enteritis model. Related to Figure 5.**

(A) Representative H&E staining of the small intestine at day 3 after 5-FU treatment. The tissue was sectioned in a Swiss-roll configuration, with the proximal region oriented at the center and the distal region at the periphery. Boxed regions are enlarged to the right. H&E-stained sections from vehicle-treated mice are also shown below for comparison.

(B) Body weight curve of mice in the 5-FU enteritis model. Data are presented as mean ± SD (n = 12).

(C) Quantification of clonal ribbon formation within 5 cm distal to the pylorus. Data are presented as mean ± SD (n = 3 per group, two-tailed Student's *t*-test, $^{*}P < 0.05$).
